# Non-invasive detection of upper tract urothelial carcinomas through the analysis of driver gene mutations and aneuploidy in urine

**DOI:** 10.1101/203968

**Authors:** Simeon Springer, Chung-Hsin Chen, Lu Li, Chris Douville, Yuxuan Wang, Josh Cohen, Bahman Afsari, Ludmila Danilova, Natalie Silliman, Joy Schaeffer, Janine Ptak, Lisa Dobbyn, Maria Papoli, Isaac Kinde, Chao-Yuan Huang, Chia-Tung Shun, Rachel Karchin, George J. Netto, Robert J. Turesky, Byeong Hwa Yun, Thomas A. Rosenquist, Ralph H. Hruban, Yeong-Shiau Pu, Arthur P. Grollman, Cristian Tomasetti, Nickolas Papadopoulos, Kenneth W. Kinzler, Bert Vogelstein, Kathleen G. Dickman

## Abstract

Upper tract urothelial carcinomas (UTUC) of the renal pelvis or ureter can be difficult to detect and challenging to diagnose. Here, we report the development and application of a non-invasive test for UTUC based on molecular analyses of DNA recovered from cells shed into the urine. The test, called UroSEEK, incorporates assays for mutations in eleven genes frequently mutated in urologic malignancies and for allelic imbalances on 39 chromosome arms. At least one genetic abnormality was detected in 75% of urinary cell samples from 56 UTUC patients but in only 0.5% of 188 samples from healthy individuals. The assay was considerably more sensitive than urine cytology, the current standard-of-care. UroSEEK therefore has the potential to be used for screening or to aid in diagnosis in patients at increased risk for UTUC, such as those exposed to herbal remedies containing the carcinogen aristolochic acid.

## Introduction

More than 400,000 new cases of urologic transitional cell carcinoma are diagnosed worldwide each year (Antoni et al., 2017). Although most of these urothelial carcinomas arise in the bladder in the lower urinary tract, 5-10% originate in the upper urinary tract in the renal pelvis and/or ureter (Roupret et al., 2015; Soria et al., 2017). The annual incidence of these upper tract urothelial carcinomas (UTUC) in Western countries is 1-2 cases per 100,000 (Roupret et al., 2015; Soria et al., 2017), but occurs at a much higher rate in populations exposed to aristolochic acid (AA) (Chen et al., 2012; Grollman, 2013; Lai et al., 2010; Taiwan Cancer Registry, 2017). AA is a carcinogenic and nephrotoxic nitrophenanthrene carboxylic acid produced by *Aristolochia* plants (Hsieh et al., 2008; National Toxicology Program, 2011). An etiological link between AA exposure and UTUC has been established in two distinct populations. The first resides in Balkan countries where *Aristolochia plants* grow naturally in wheat fields (Jelakovic et al., 2012). The second population is in Asia, where *Aristolochia* herbs are widely used in the practice of Traditional Chinese Medicine (Grollman, 2013; National Toxicology Program, 2011). The public health threat posed by the medicinal use of *Aristolochia* herbs is exemplified by Taiwan, which has the highest incidence of UTUC in the world (Chen et al., 2012; Yang et al., 2002). More than one-third of the adult population in Taiwan has been prescribed herbal remedies containing AA (Hsieh et al., 2008), resulting in an unusually high (37%) proportion of UTUC cases relative to all urothelial cancers (Taiwan Cancer Registry, 2017).

Nephroureterectomy can be curative for patients with UTUC when it is detected at an early stage (Li et al., 2008). However, these cancers are largely silent until the onset of overt clinical symptoms, typically hematuria, and as a result, most patients are diagnosed only at an advanced stage (Roupret et al., 2015). Diagnostic tests for the detection of early-stage UTUC are not currently available. There is thus a need for clinical tools that can be used to identify early UTUCs in populations at risk for developing this type of malignancy. Relapse following surgery is also a concern, as UTUC can recur in the contralateral upper urinary tract and/or in the bladder (Roupret et al., 2015; Soria et al., 2017). Vigilant surveillance for signs of malignancy is therefore an essential part of follow-up care in UTUC patients, and non-invasive tests for recurrent disease could substantially improve post-surgical management, particularly as urine cytology cannot detect the majority of UTUCs (Baard et al., 2017).

As UTUCs are in direct contact with the urine, we hypothesized that genetic analyses of exfoliated urinary cells could be used to detect upper urinary tract neoplasm in a noninvasive fashion (Figure 1). In the current study, we tested this hypothesis through the analysis of urinary cell DNA using assays that could identify a variety of genetic abnormalities.

**Figure 1.**
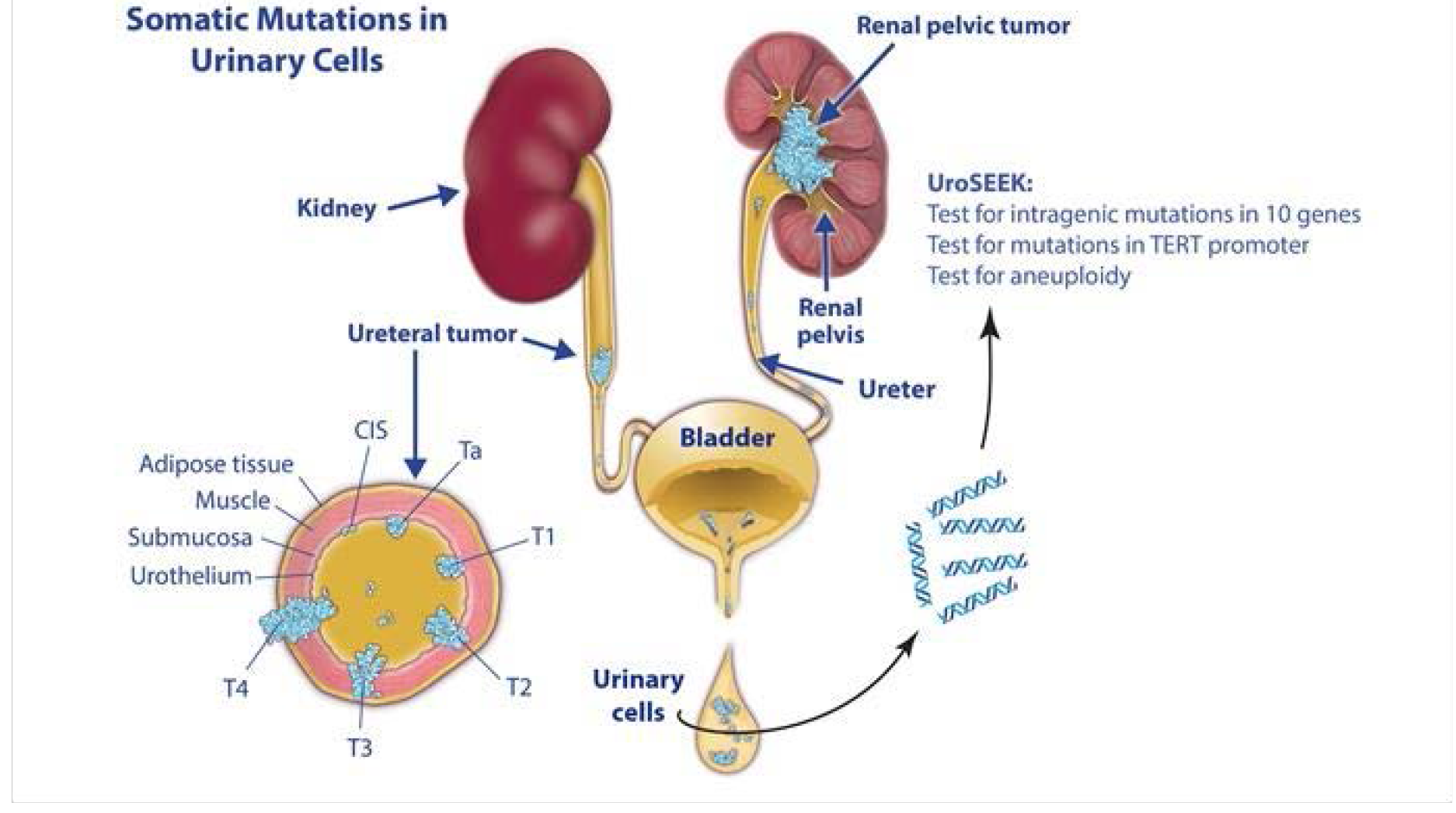
Non-invasive detection of upper tract urothelial cancer (UTUC) through genetic analysis of urinary cell DNA. Upper urinary tract tumors arise in the renal pelvis and/or ureter and are in direct contact with urine Urine contains a mixture of normal cells that are constitutively shed from various sites along the urinary system, along with malignant cells when present (blue cells in figure) The UroSEEK assay relies on mutational analyses of genes frequently mutated in urinary cancers along with a determination of chromosome losses and gains.

## Results

<u>Cohort characteristics.</u> Thirty-two females and twenty-four males ranging in age from 39-85 years participated in the study (summary in Table 1; individual data are in Supplementary File 1). This gender distribution, atypical of UTUC patients in Western countries where males predominate (Shariat et al., 2011), is consistent with previous epidemiologic studies of Taiwanese individuals with known exposures to AA (Chen et al., 2012). Tobacco use was reported by 18% of this cohort, all males. Based on estimated glomerular filtration rate (eGFR) values, renal function was unimpaired (chronic kidney disease (CKD) stage 0-2) in 45% of the subjects, while mild-to-moderate renal disease (CKD stage 3) or severe disease (CKD stages 4-5) was noted for 43% and 12% of the cohort, respectively (Table 1).

**Table 1.**
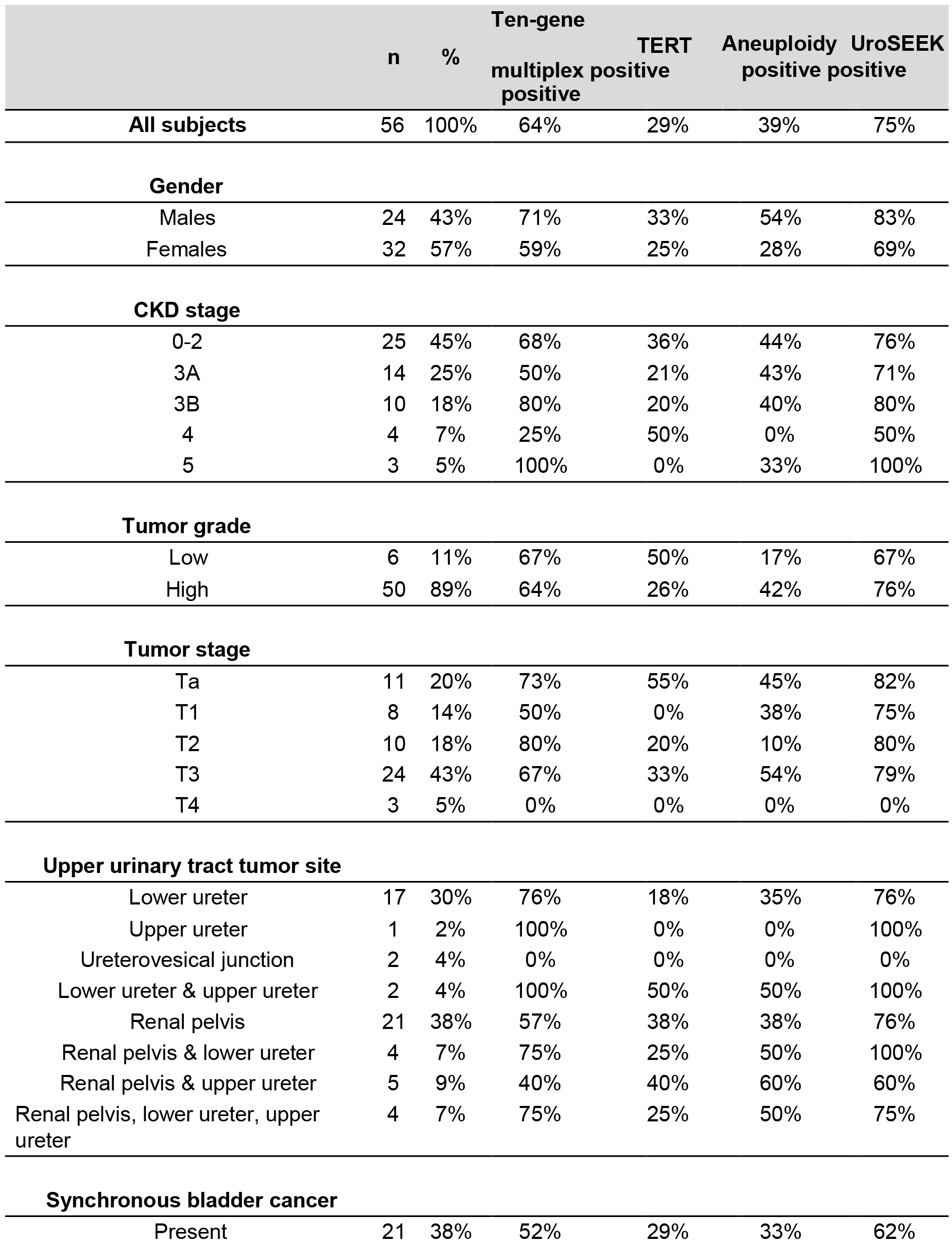
**Demographic, clinical and genetic features of the UTUC cohort stratified by UroSEEK results.**

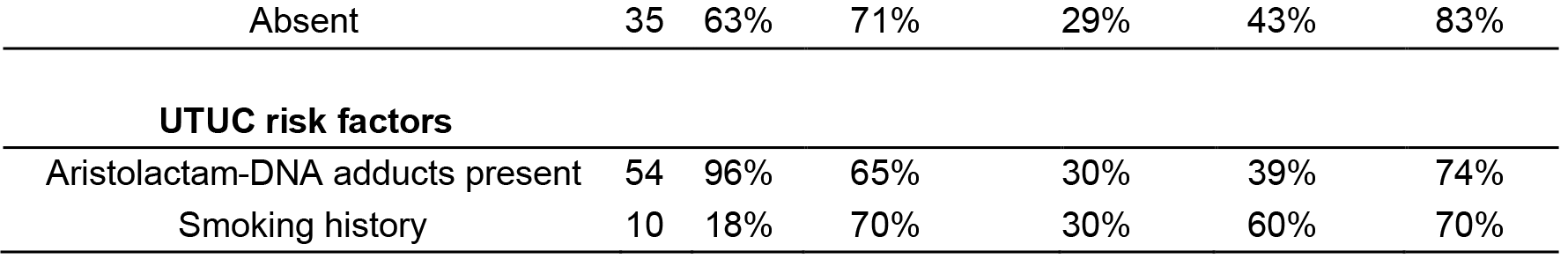

Tumors were confined to a single site along the upper urinary tract in the majority of cases (38% renal pelvis; 39% ureter), while multifocal tumors affecting both renal pelvis and ureter occurred in 23% of the patients. Synchronous bladder cancer (diagnosed within 3 months prior to nephroureterectomy) was present in 38%. Histologically, 89% of the tumors were classified as high grade, with the majority categorized as muscle- invasive (T2-T4, 66%) (Table 1).

<u>Mutational analysis.</u> We performed three separate tests for genetic abnormalities that might be found in urinary cells derived from UTUCs (Figure 2, Supplementary Files 2-5). First, we evaluated mutations in selected exomic regions of ten genes *(CDKN2A*, *ERBB2*, *FGFR3*, *HRAS*, *KRAS*, *MET*, *MLL*, *PIK3CA*, *TP53*, and *VHL*) that are frequently altered in urologic tumors (Sfakianos et al., 2015). For this purpose, we designed a specific set of multiplex primers that allowed us to detect mutations in as few as 0.03% of urinary cells (Supplementary File 6). The capacity to detect such low mutant fractions was a result of the incorporation of molecular barcodes in each of the primers, thereby substantially reducing the artifacts associated with massively parallel sequencing (Kinde et al, 2011).

**Figure 2.**
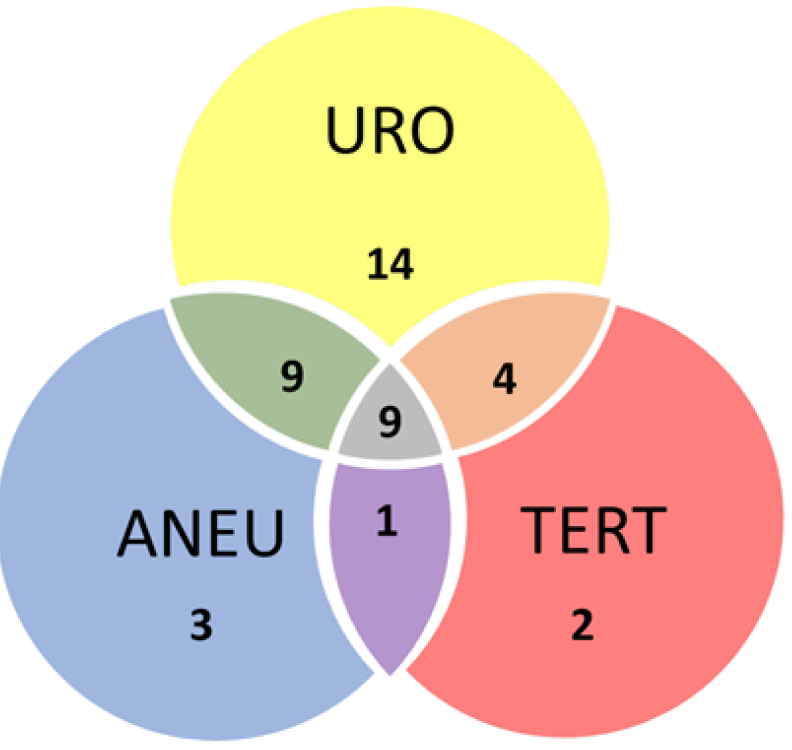
Venn diagram showing the distribution of positive results for each of the three UroSEEK assays.

Second, we evaluated *TERT* promoter mutations, based on prior evidence that *TERT* promoter mutations are often found in UTUCs (Kinde et al., 2013). A singleplex PCR was used for this analysis because the unusually high GC-content of the *TERT* promoter precluded its inclusion in the multiplex PCR design. Third, we evaluated the extent of aneuploidy using a technique in which a single PCR is used to co-amplify ~38,000 members of a subfamily of long interspersed nucleotide element-1 (L1 retrotransposons) (Kinde et al., 2012). L1 retrotransposons, like other human repeats, have spread throughout the genome via retrotransposition and are found on all 39 non-acrocentric autosomal arms (Ostertag & Kazazian, 2001).

The multiplex assay detected mutations in 36 of the 56 urinary cell samples from UTUC patients (64%, 95% CI 51 % to 76% (Table 1 and Supplementary File 2)). A total of 57 mutations were detected in nine of the ten target genes (Figure 3). The median mutant allele frequency (MAF) in the urinary cells was 5.6% and ranged from 0.3% to 80%. The most commonly altered genes were *TP53* (58% of the 57 mutations) and *FGFR3* (16% of the 57 mutations) (Figure 3). The distribution of mutant genes was roughly consistent with expectations based on previous exome-wide sequencing studies of UTUCs (Moss et al., 2017). None of the 188 urinary cell samples from healthy individuals had a detectable mutation in any of the ten genes assayed (100% specificity, CI 97.5% to 100%).

**Figure 3.**
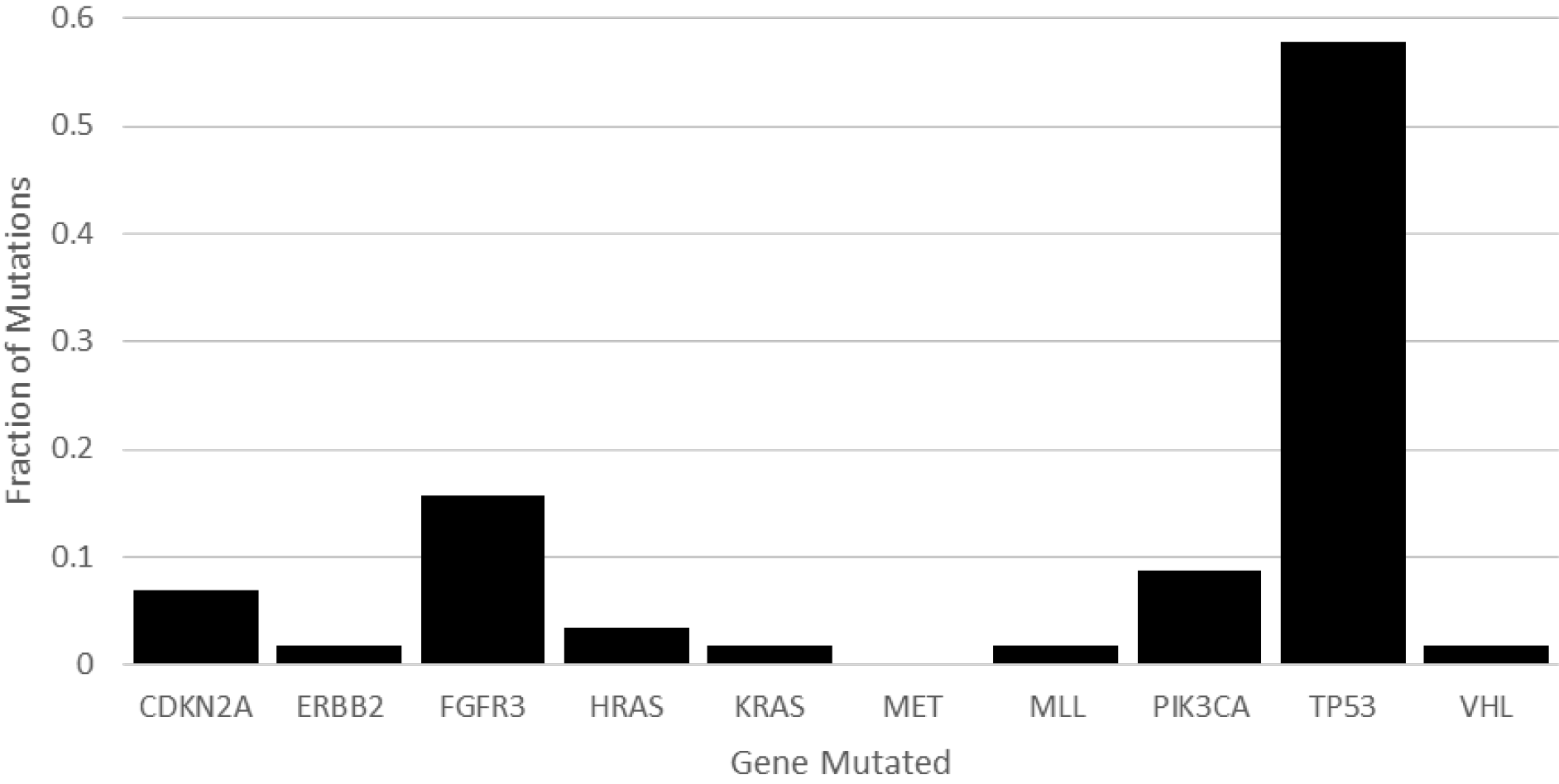
Fraction of total mutations for each gene in the 10-gene panel used to analyze urinary cell DNA from UTUC patients.

Mutations in the *TERT* promoter were detected in 16 of the 56 urinary cell samples from UTUC patients (29%, 95% CI 18% to 42%) (Table 1 and Supplementary File 3). The median *TERT* MAF in the urinary cells was 2.22% and ranged from 0.59% to 46.3%. One of the 188 urinary samples from healthy individuals harbored a mutation *(TERT* g.1295250C>T with a MAF of 0.39%). In the UTUC urinary cell samples, mutations were detected in three positions: 94% of the mutations were at hg1295228 (67%) and hg1295250 (28%), which are 69 and 91 bp upstream of the transcription start site, respectively. These positions have been previously shown to be critical for the appropriate transcriptional regulation of *TERT*. In particular, the mutant alleles recruit the GABPA/B1 transcription factor, resulting in the H3K4me2/3 mark of active chromatin and reversing the epigenetic silencing present in normal cells (Stern et al., 2015).

<u>Aristolochic acid exposure.</u> The activated metabolites of aristolochic acid bind covalently to the exocyclic amino groups in purine bases, with a preference for dA, leading to characteristic A>T transversions (Hollstein et al., 2013). To determine whether the individuals in our cohort had been exposed to AA, we quantified renal cortical DNA adducts using mass spectrometry (Yun et al., 2012). All but two of the 56 patients had detectable aristolactam (AL)-DNA adducts (Table 1) with levels ranging from 0.4 to 68 dA-AL adducts per 10^8^ nucleotides. Moreover, the A>T signature mutation (Hoang et al., 2016) associated with AA was highly represented in the mutational spectra of *TP53* (18/32 A>T) and *HRAS* (2/2 A>T) found in urinary cells (Supplementary File 3).

<u>Aneuploidy.</u> Aneuploidy was detected in 22 of the 56 urinary cell samples from UTUC patients (39%, 95% CI 28% to 52%, Supplementary Files 4 and 5) but in none of the 188 urinary cell samples from healthy individuals. The most commonly altered arms were 1q, 7q, 8q, 17p, and 18q. Some of these arms harbor well-known tumor oncogenes or suppressor genes that have been shown to undergo changes in copy numbers in many cancers (Vogelstein et al., 2013).

<u>Comparison with primary tumors.</u> Tumor samples from all 56 patients enrolled in this study were available for comparison and were studied with the same three assays used to analyze the urinary cell samples. This comparison served two purposes. First, it allowed us to determine if the mutations identified in the urinary cells were derived from the available tumor specimen from the same patient. There were a total of 39 UTUC cases in which a mutation could be identified in the urinary cells. In 35 (90%) of these 39 cases, at least one of the mutations identified in the urine sample (Supplementary Files 2 and 3) was also identified in the corresponding tumor DNA sample (Supplementary Files 6 and 7). When all 80 mutations identified in the urinary cells were considered, 63 (79%) were identified in the corresponding tumor sample (Supplementary Files 6 and 7). In any of the three assays, the discrepancies between urine and tumor samples might be explained by the fact that we had access to only one tumor per patient, even though more than one anatomically distinct tumor was often evident clinically (Table 1). Additionally, DNA was extracted from only one piece of tissue from each tumor, and intratumoral heterogeneity (McGranahan et al., 2015) could have been responsible for some of the discrepancies.

The tumor data helped determine why 17 of the 56 urinary cell samples from UTUC patients did not contain detectable mutations. The reason could either have been that the primary tumors did not harbor a mutation present in our gene panel or that the primary tumor did contain such a mutation but the fraction of neoplastic cells in the urine sample was not high enough to allow its detection. From the evaluation of the primary tumor samples, we found that four (24%) of the 17 urine samples without detectable mutations were from patients whose tumors did not contain any of the queried mutations (Supplementary Files 6 and 7). We conclude that the main reason for failure of the mutation test was an insufficient number of cancer cells in the urine, and this accounted for 13 (76%) of the 17 failures. There were 22 cases in which aneuploidy was observed in the urinary cell samples. Overall, 96% of the chromosome gains or losses observed in the urinary cells were also observed in the primary tumors (examples in Figure 4). Conversely there were 34 cases in which aneuploidy was *not* observed in the urinary cell samples. Evaluation of the 56 tumors with the same assay showed that all but three were aneuploid, so as with mutations, the main reason for failure of the aneuploidy assay was insufficient amounts of neoplastic DNA in the urinary cells.

There were 22 cases in which aneuploidy was observed in the urinary cell samples. Overall, 96% of the chromosome gains or losses observed in the urinary cells were also observed in the primary tumors (examples in Figure 4). Conversely there were 34 cases in which aneuploidy was not observed in the urinary cell samples. Evaluation of the 56 tumors with the same assay showed that all but three were aneuploid, so as with mutations, the main reason for failure of the aneuploidy assay was insufficient amounts of neoplastic DNA in the urinary cells.

**Figure 4.**
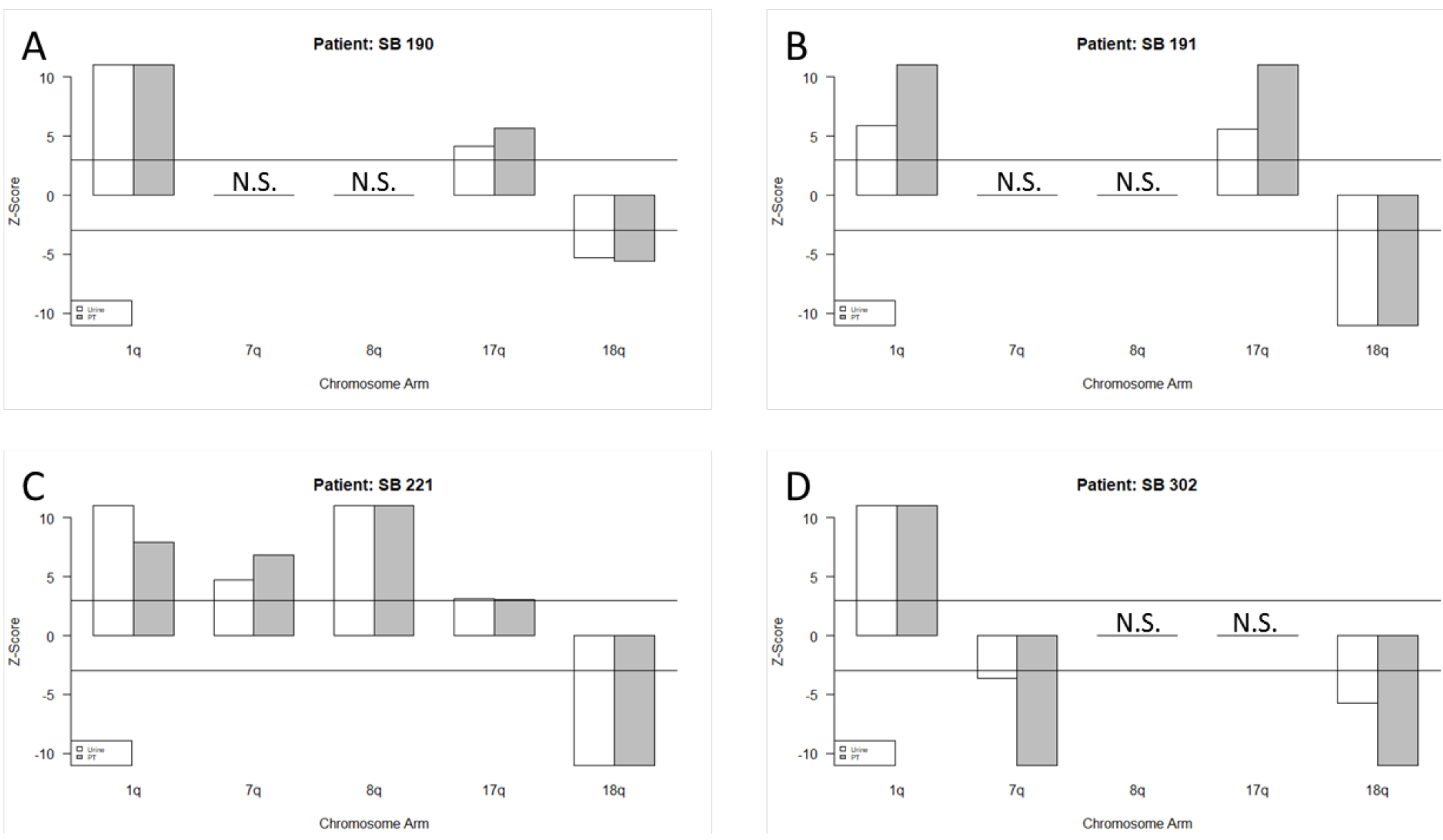
Comparison of copy number variations in matched tumor and urinary cell DNA samples from four individual UTUC patients. Z-scores >3 or <-3 were considered as significant for chromosome gains or losses, respectively. N.S., not significant. Data for all 56 patients are provided in Supplementary File 5.

<u>Biomarkers in combination.</u> There are two factors that limit sensitivity for genetically based biomarkers. First, a sample can only be scored as positive for the biomarker if it contains DNA from a sufficient number of neoplastic cells to be detected by the assay. Second, the tumor from which the neoplastic cells were derived must harbor the genetic alteration that is queried. Combination assays can increase sensitivity by assessing more genetic alterations, and are thereby more likely to detect at least one genetic alteration present in the tumor. However, mutations in clinical samples often are present at low allele frequencies (Supplementary Files 2 and 3), requiring high coverage of every base queried. It would be prohibitively expensive to perform whole exome sequencing at 10,000× coverage, for example, so some compromise is needed. In our study, we evaluated carefully selected regions of 11 genes (including *TERT*) together with copy number analysis of 39 chromosome arms. Even if a tumor does not contain a genetic alteration in one of the 11 genes assessed, it might still be aneuploid and detectable by the urinary cell assay for aneuploidy. The sensitivity of aneuploidy detection is less than that of the mutation assays. Simulations showed that DNA containing a minimum of 1% neoplastic cells is required for reliable aneuploidy detection, while mutations present in as few as 0.03% of the DNA templates can be detected by the mutation assays used in our study (Kinde et al 2013; Bettegowda et al. 2014, Wang et al., 2016). Nevertheless, urinary cell samples that had relatively high fractions of neoplastic cells but did not contain a detectable mutation in the 11 queried genes should still be detectable by virtue of their aneuploidy because, as noted above, 53/56 UTUCs studied here were aneuploid. Additionally, some of the mutations in the 11 genes queried, such as large insertions or deletions or complex changes, might be undetectable by mutation-based assays but a sample with such an undetectable mutation could still score positive in a test for aneuploidy.

To determine whether these theoretical arguments made a difference in practice, we evaluated biomarker performance with the combined approaches, collectively called UroSEEK. As noted above, the ten-gene multiplex assay, the *TERT* singleplex assay, and the aneuploidy assays yielded 64, 29%, and 39% sensitivities, respectively, when used separately (Table 1). Twenty-three samples without TERT promoter mutations tested positive for mutations in one of the other ten genes (Venn diagram in Figure 2). Conversely, three samples without detectable mutations with the multiplex assay scored positive for *TERT* promoter mutations (Figure 2). And, three of the urinary cell samples without any detectable mutations were positive for aneuploidy (Figure 2). Thus, when the three assays were used together, and a positive result in any one assay was sufficient to score a sample as positive, the sensitivity rose to 75% (95% CI 62.2% to 84.6%). Only one of the 188 samples from healthy individuals scored positive in the UroSEEK test (specificity 99.5%, CI 97.5 to 100%).

To determine the basis for the increased sensitivity afforded by the combination assays, we evaluated data from the primary tumors of the three patients whose urinary cell samples exhibited aneuploidy but did not harbor detectable mutations. We found that these three tumors did not contain any mutations in the 11 queried genes, explaining why these same assays were negative when applied to urinary cell DNA. As noted above, these three tumors were aneuploid, thus affording the opportunity to detect these copy number variations in the urinary cell samples.

<u>Correlation with clinical features.</u> One of the most important properties of a cancer biomarker is that it be able to detect tumors at an early stage, enabling surgical removal of the lesions prior to widespread metastasis. Fortunately, UroSEEK appeared to be as sensitive for detecting both early and late tumors. It scored positive in 15 (79%) of 19 patients with stage Ta or T1 tumors and in 27 (73%) of 37 patients with stage T2-T4 tumors (Table 1). Ten-year cancer specific survival rates show that 91% of UTUC patients with stage T1 malignancies are expected to be cured by surgery, compared to only 78%, 34% and 0% of patients with stage 2, 3, or 4 tumors, respectively (Li et al., 2008).

UroSEEK sensitivity was independent of a variety of clinical parameters other than tumor stage, including gender, CKD stage, tumor grade, tumor location and risk factors for developing UTUC (Table 1), indicating that the assay should be suitable for evaluation of diverse patient populations. Furthermore, UroSEEK was considerably more sensitive than urine cytology in this cohort. Cytology was available in 42 cases, and of these only four (9.5%) were diagnosed as carcinoma cytologically. Even if samples scored as “suspicious for malignancy” by cytology were considered as positive, the sensitivity was only 26% (including the four scored as positive and seven scored as suspicious). UroSEEK detected all four cases scored as positive by cytology, five of the seven cases scored as suspicious for malignancy, and 22 of the 31 samples scored by cytology as inconclusive or negative.

## Discussion

The National Health Insurance database indicates that one-third of the adult population in Taiwan had been prescribed *Aristolochia* herbs between 1997 and 2003 (Hsieh et al., 2008). Additional people in Taiwan are exposed to *Aristolochia* herbs through nonprescribed, herbal medicines. It has been estimated that 100 million people in China are at risk for UTUC as a result of exposure to this carcinogen (Grollman, 2013; Hu et al., 2004). Non-invasive methods to screen the large numbers of at-risk individuals for UTUC in such populations are thus clearly desirable. Currently, no such screening methods are available. Urine cytology requires highly trained individuals and even in expert hands is not particularly sensitive for urothelial carcinoma (Lotan & Roehrborn, 2003; Netto & Tafe, 2016; Zhang et al., 2016). In addition, although urine cytology has value for the detection of high-grade neoplasms, it is unable to detect the vast majority of low-grade tumors. A large number of tests yield ‘atypical cells”, which are uninterpretable with respect to malignancy (Barkan et al., 2016). Radiologic tests, such MRI or CT-scans, are not well suited for screening and the latter confers significant radiation exposure. Ureteroscopy is often definitive, but in addition to being invasive, requires highly skilled clinicians and is also ill-suited as a screening tool (Golan et al., 2015).

Liquid biopsy has recently gained attention as a non-invasive approach for cancer detection. Although this concept often refers to blood samples, it can be applied to other body fluids, such as urine (Patel et al., 2017; Sidransky et al., 1991; Togneri et al., 2016), which contains DNA from several sources including (i) glomerular filtration of circulating free DNA (Botezatu et al., 2000) released by normal and tumor cells from sites throughout the body; (ii) DNA released directly into urine by normal and tumor cells of the urinary tract; and (iii) intact normal or malignant cells of the urinary tract exfoliated into urine (Bettegowda et al., 2014; Dawson et al., 2013; Dressman et al., 2003; Forshew et al., 2012; Haber & Velculescu, 2014; Kinde, Bettegowda, et al., 2013; Springer et al., 2015; Vogelstein & Kinzler, 1999; Wang, Springer, Mulvey, et al., 2015; Wang, Springer, Zhang, et al., 2015; Wang et al., 2016). We chose the latter option for the current study to increase the sensitivity and specificity of our biomarker assay.

While optimizing conditions for the current study, we compared the relative performance of mutation assays in matched plasma and urine samples obtained from 14 UTUC patients. In each case, a *TERT* or *TP53* mutation was first identified in the primary tumor, then that particular mutation was queried in DNA from the urine or plasma using a singleplex assay. Mutations were detected in 93% (13/14) of the urinary cell DNA samples compared to 36% (5/14) of the plasma samples. Importantly, the plasma test failed to identify any of the six non-muscle-invasive cancers (Ta/T1), while all six (100%) were identified in the matched urinary cell DNA samples. The superior performance in urinary cells was likely due to a substantial enrichment for mutated DNA in these cells compared to plasma: the median MAF in plasma when a mutation was detectable was only 0.3% compared to 15% in the urinary cells.

Although the approach described here has significant potential for screening purposes, we emphasize that the current study demonstrates proof-of-principle rather than clinical applicability. Accordingly, there are several caveats to the study that are worthy of attention. First, we only evaluated 56 patients. Second, the study was in essence retrospective rather than prospective. Another “caveat” is that our assays on urinary cells cannot distinguish between UTUC and bladder tumors, and the UroSEEK test could in theory also detect kidney cancer. We consider this a strength rather than a weakness, because the detection of bladder or kidney cancer is as important as the detection of UTUC. Bladder cancers are more common than UTUC in the general population (Roupret et al., 2015), and patients exposed to AA are at risk for bladder cancer and renal cell carcinoma as well as for UTUC (Hoang et al., 2016; Lai et al., 2010).

The current study establishes the conceptual and molecular biological foundation for two clinical trials that are now being planned. The first is a prospective screening study of urinary cells obtained from individuals in Taiwan with microscopic hematuria, many of whom are at risk of developing UTUC due to past exposure to AA. The purpose of this study will be to determine whether UTUC or bladder cancer can be detected by UroSEEK prior to the advent of hematuria. The second is a prospective study of patients with UTUCs that have been surgically removed. Recurrence of disease in the bladder occurs in 22 to 47% of UTUC cases, presumably from cells seeded from the upper urinary tract, while tumors in the contralateral tract appear in an additional 2 to 6% of patients (Roupret et al., 2015). The purpose of this trial is to determine whether the analysis of urinary cell DNA can reveal recurrent or new disease earlier than conventional clinical, cytologic, or radiologic methods.

## Methods

<u>Cohort studied.</u> Sequential patients with UTUC scheduled to undergo a radical unilateral nephroureterectomy at National Taiwan University Hospital in 2012-2016 were asked to participate in the study. All patients provided informed consent using the consent form and study design reviewed and approved by the Institutional Review Boards at National Taiwan University and Stony Brook University. A total of 56 UTUC patients were enrolled in the study after excluding four patients with gross hematuria and one patient with a tumor-urine DNA mismatch by identity testing (see below). Urinary cell DNA from 188 urine samples donated by healthy individuals in the U.S. of average age 40, range 19 to 60 years old, was used to assess the specificity of the UroSEEK test. White blood cell (WBC) DNA from 94 normal individuals from the U.S. was used to evaluate the technical specificity of the PCR analysis.

<u>Biological samples.</u> Patients provided urine samples one day prior to surgery. Urinary cells were isolated by centrifugation at 581*g* for 10 min at room temperature, washed thrice in saline using the same centrifugation conditions, and stored frozen until DNA was isolated using a Qiagen kit #937255 (Germantown, MD). DNA was purified from fresh-frozen resected samples of upper tract tumors and renal cortex by standard phenol-chloroform extraction procedures (Chen et al., 2012; Jelakovic et al., 2012). One upper urinary tract tumor per patient was analyzed; for cases with tumors at multiple sites, renal pelvic tumors were preferentially selected whenever available. Formalinfixed, paraffin-embedded tumor samples were staged and graded by a urologic pathologist, and the presence of one or more upper tract urothelial carcinomas was confirmed by histopathology for each enrolled subject. Pertinent clinical and demographic data were obtained by a chart review of each subject. eGFR was calculated by the MDRD equation (Levey et al., 2006) and used to determine CKD stage(Levey et al., 2005).

<u>DNA adduct analysis.</u> AL-DNA adduct (7-(deoxyadenosin-*N*^6^-yl) aristolactam I; dA-AL-I) levels in 2 ug of DNA from the normal renal cortex of UTUC patients were quantified by ultra-performance liquid chromatography-electrospray ionization/multistage mass spectrometry (UPLC-ESI/MS^n^) with a linear quadrupole ion trap mass spectrometer (LTQ Velos Pro, Thermo Fisher Scientific, San Jose, CA) as described previously(Yun et al., 2012).

<u>Mutation analysis</u>. Three separate assays were used to search for abnormalities in urinary cell DNA. First, a multiplex PCR was used to detect mutations in regions of ten genes commonly mutated in urologic malignancies *CDKN2A*, *ERBB2*, *FGFR3*, *HRAS*, *KRAS*, *MET*, *MLL*, *PIK3CA*, *TP53*, and *VHL* (Cancer Genome Atlas Research, 2014; Lin et al., 2010; Mo et al., 2007; Netto, 2011; Sarkis et al., 1995; Sarkis et al., 1994; Sarkis et al., 1993; Wu, 2005). The 57 primer pairs used for this multiplex PCR were divided in a total of three multiplex reactions, each containing non-overlapping amplicons (Supplementary File 6) These primers were used to amplify DNA in 25 uL reactions as previously described (Kinde et al., 2011) except that 15 cycles were used for the initial amplification. Second, the *TERT* promoter region was evaluated. A single amplification primer pair was used to amplify a 73-bp segment containing the region of the *TERT* promoter known to harbor mutations in BC (Kinde, Munari, et al., 2013). The conditions used to amplify it were the same as used in the multiplex reactions described above except that Phusion GC Buffer (Thermo-Fisher) instead of HF buffer was used and 20 cycles were used for the initial amplification. Note that the *TERT* promoter region could not be included in the multiplex PCR because of the high GC content of the former. PCR products were purified with AMPure XP beads (Beckman Coulter, PA, USA) and 0.25% of the purified PCR products (multiplex) or 0.0125% of the PCR products (*TERT* singleplex) were then amplified in a second round of PCR, as described in (Wang et al. 2015). The PCR products from the second round of amplification were then purified with AMPure and sequenced on an Illumina instrument. For each mutation identified, the mutant allele frequency (MAF) was determined by dividing the number of uniquely identified reads with mutations (Kinde et al., 2011) by the total number of uniquely identified reads. Each DNA sample was assessed in two independent PCRs, for both the *TERT* promoter and multiplex assays, and samples were scored as positive only if both PCRs showed the same mutation. The mutant allele frequencies and number of unique templates analyzed listed in the Supplementary Files refer to the average of the two independent assays.

To evaluate the statistical significance of putative mutations, we assessed DNA from WBCs of 188 unrelated normal individuals. A variant observed in the samples from cancer patients was only scored as a mutation if it was observed at a much higher MAF than observed in the normal WBCs used as controls. Specifically, the classification of a sample's DNA status was based on two complementary criteria applied to each mutation: 1) the difference between the average MAF in the sample of interest and the corresponding maximum MAF observed for that same mutation in a set of controls, and 2) the Stouffer’s Z-score obtained by comparing the MAF in the sample of interest to a distribution of normal controls. To calculate the Z-score, the MAF in the sample of interest was first normalized based on the mutation-specific distributions of MAFs observed among all controls. Following this mutation-specific normalization, a P-value was obtained by comparing the MAF of each mutation in each well with a reference distribution of MAFs built from normal controls where all mutations were included. The Stouffer's Z-score was then calculated from the p-values of two wells, weighted by their number of UIDs. The sample was classified as positive if either the difference or the Stouffer’s Z-score of its mutations was above the thresholds determined from the normal WBCs. The threshold for the difference parameter was defined by the highest MAF observed in any normal WBCs. The threshold for the Stouffer′s Z-score was chosen to allow one false positive among the 188 normal urine samples studied.

<u>Analysis of aneuploidy</u>. Aneuploidy was assessed with FastSeqS, which uses a single primer pair to amplify ~38,000 loci scattered throughout the genome (Kinde et al., 2012). After massively parallel sequencing, gains or losses of each of the 39 chromosome arms covered by the assay were determined using a bespoke statistical learning method described elsewhere (Deauville et al., in preparation). A support vector machine (SVM) was used to discriminate between aneuploid and euploid samples. The SVM was trained using 3150 low neoplastic cell fraction synthetic aneuploid samples and 677 euploid peripheral white blood cell (WBC) samples. Samples were scored as positive when the genome-wide aneuploidy score was >0.7 and there was at least one gain or loss of a chromosome arm.

<u>Identity checks</u>. A multiplex reaction containing 26 primers detecting 31 common SNPs on chromosomes 10 and 20 was performed using the amplification conditions described above for the multiplex PCR. The 26 primers used for this identity evaluation are listed in Supplementary File 9.

<u>Statistical analysis.</u> Performance characteristics of urine cytology, UroSEEK and its three components were calculated using using MedCalc statistical software, online version (https://www.medcalc.org/calc/diagnostic_test.php). Confidence intervals (95%) were determined with an online GraphPad Software Inc. statistical calculator (https://www.graphpad.com/quickcalcs/confInterval1/) using the modified Wald method.

## Acknowledgements

This research was supported, in part, by a grant (104-2314-B-002 -132-) to CHC from the National Science Council, Taiwan, and by a grant (104-2314-B-002-121-MY3) to YSP from the National Science Council, Taiwan. We appreciate the clinical services provided by Dr. Kuo-How Huang, Shuo-Meng Wang, Huai-Ching Tai and Yuan-Ju Lee (Department of Urology, National Taiwan University Hospital). The generous support provided by Henry and Marsha Laufer is gratefully acknowledged, as is support from the Virginia and D.K. Ludwig Fund for Cancer Research and the Conrad R. Hilton Foundation. All sequencing was performed at the Sol Goldman Sequencing Facility at Johns Hopkins. This work was also supported by grants from the National Institute of Environmental Health Sciences (R01 ES019564) and the National Institutes of Health (Grants CA 77598, CA 06973, and GM 07309). The authors appreciate the medical illustrations skillfully designed by Kathleen Gebhart.

